# Imaging demyelinated axons after spinal cord injuries with PET tracer [^18^F]3F4AP

**DOI:** 10.1101/2024.04.24.590984

**Authors:** Karla M. Ramos-Torres, Sara Conti, Yu-Peng Zhou, Amal Tiss, Celine Caravagna, Kazue Takahashi, Zhigang He, Moses Q. Wilks, Sophie Eckl, Yang Sun, Jason Biundo, Kuang Gong, Zhigang He, Clas Linnman, Pedro Brugarolas

## Abstract

Spinal cord injuries (SCI) often lead to lifelong disability. Among the various types of injuries, incomplete and discomplete injuries, where some axons remain intact, offer potential for recovery. However, demyelination of these spared axons can worsen disability. Demyelination is a reversible phenomenon, and drugs like 4-aminopyridine (4AP), which target K^+^ channels in demyelinated axons, show that conduction can be restored. Yet, accurately assessing and monitoring demyelination post-SCI remains challenging due to the lack of suitable imaging methods. In this study, we introduce a novel approach utilizing the positron emission tomography (PET) tracer, [^18^F]3F4AP, specifically targeting K^+^ channels in demyelinated axons for SCI imaging. Rats with incomplete contusion injuries were imaged up to one month post-injury, revealing [^18^F]3F4AP’s exceptional sensitivity to injury and its ability to detect temporal changes. Further validation through autoradiography and immunohistochemistry confirmed [^18^F]3F4AP’s targeting of demyelinated axons. In a proof-of-concept study involving human subjects, [^18^F]3F4AP differentiated between a severe and a largely recovered incomplete injury, indicating axonal loss and demyelination, respectively. Moreover, alterations in tracer delivery were evident on dynamic PET images, suggestive of differences in spinal cord blood flow between the injuries. In conclusion, [^18^F]3F4AP demonstrates efficacy in detecting incomplete SCI in both animal models and humans. The potential for monitoring post-SCI demyelination changes and response to therapy underscores the utility of [^18^F]3F4AP in advancing our understanding and management of spinal cord injuries.

## Introduction

Spinal cord injury (SCI) can result in devastating loss of mobility, often causing life-long struggles to regain independence and quality of life. During the initial months following injury, there are rapid degenerative changes, such as demyelination at and around the lesion, which are associated with a less favorable recovery outcome(*1–3*). Demyelination, which causes slowed or failed axonal conduction, is thought to be a major contributor to disability after incomplete SCI(*2, 4*). Furthermore, with the loss of myelin, spared demyelinated axons are directly exposed to inflammatory cytokines and free radicals, potentially leading to further neuronal loss via necrosis or apoptosis. Consequently, therapeutic approaches that promote remyelination are a major research focus in SCI, with much knowledge gained from multiple sclerosis (MS). These include pharmacological-, cell-, physical-, and electrical stimulation-therapies(*4–16*). Although many of these approaches have shown promise in standardized animal models, a significant challenge in translating these treatments to humans is determining how to monitor their response, considering the heterogenous nature of human SCI. Existing magnetic resonance imaging (MRI) methods for following SCI progression rely on anatomical changes that reflect the ultimate outcome of the demyelination or remyelination process(*17*). These approaches lack specificity and exhibit only modest correlation with clinical symptoms. In addition, the presence of spinal cord stabilization hardware can create magnetic resonance artifacts, restricting the feasibility of using MRI to directly assess characteristics of the spinal cord lesion and proximal areas of the cord(*18*).

Positron emission tomography (PET) has the potential to offer sensitive and quantitative imaging of biochemical changes following SCI. Various PET tracers have been employed to evaluate the biological and physiological outcomes of SCI(*19*). Owing to its biochemical specificity, PET imaging has been used to inform on certain processes including metabolic stability as an indicator of tissue viability using [^18^F]FDG(*20*). Additionally, PET has been applied to examine the neuroinflammatory response after injury, utilizing tracers that target translocator protein 18 (TSPO) like [^18^F]GE-180(*21*). Furthermore, PET ligands can function as biomarkers and valuable tools in advancing disease monitoring and therapeutic strategies for traumatic injuries. For instance, the evaluation of axonal connectivity across a lesion through neurotransmitter-specific PET, using the presynaptic serotonin marker [^11^C]-AFM(*22*), or assessment of synaptic density loss post-injury with the selective radiotracer for synaptic vesicle glycoprotein 2A (SV2A) [^11^C]-UCB-J(*23*), can contribute to the development of innovative repair therapies.

As demyelinated neurons may contribute to disability in SCI, determining the localization and extent of demyelinated axons after injury could potentially inform disease prognosis and treatment response. Myelin-binding radiotracers based on the diamino stilbene pharmacophore have been developed to evaluate myelin changes in the spinal cord in both experimental models of demyelination(*24*) and rodent SCI models(*25, 26*). As an alternative approach, we have developed a novel PET radiotracer, 3-[^18^F]fluoro-4-aminopyridine ([^18^F]3F4AP), based on the MS drug 4-aminopyridine (4AP, dalfampridine) that targets potassium (K^+^) channels in demyelinated axons(*27*). This tracer has shown high sensitivity to demyelinated lesions in animal models of MS(*27*) and it is currently under investigation in people with MS (clinicaltrials.gov NCT04699747). Additionally, [^18^F]3F4AP has also shown high sensitivity to a minor traumatic brain injury in a rhesus macaque(*28*). Moreover, while studies in animals have shown improvement with 4AP(*29–33*), studies evaluating 4AP in people after SCI have reported mixed results(*34–42*) raising the question of who may benefit from a therapy designed to enhance conduction of demyelinated fibers. Given the sensitivity of [^18^F]3F4AP to a traumatic CNS injury, the potential therapeutic response of its parent compound, and the well-documented changes in axonal K^+^ channel expression in models of SCI(*43, 44*), we decided to investigate the use of [^18^F]3F4AP for imaging SCI. Specifically, this work examines the changes in [^18^F]3F4AP uptake in and around the injured cord in a well-established model of incomplete traumatic spinal cord injury over time. This study correlates these changes with histological alterations in myelin and the functional *in vivo* phenotype. Finally, this work provides the first images of [^18^F]3F4AP in persons with SCI, providing valuable insights into the possible clinical application of this novel PET radiotracer for monitoring and understanding SCI.

## Results

### Thoracic spinal contusion in rats as a model for SCI

Rodent models of spinal cord injury have been extensively used to examine an array of biological processes (inflammation, demyelination, axonal loss, etc.) in different degrees of injury(*45*). In this study, we chose spinal contusion in rats as a model of incomplete injury where spared demyelinated axons would be present after SCI to evaluate the ability of [^18^F]3F4AP to detect the presence of these fibers via PET imaging. Previous studies with this model have shown acute demyelination at the injury starting 7 days post injury (dpi) followed by slow remyelination(*46*). Additional studies in a related SCI compression model have shown large increases and redistribution of K_v_1.1 and K_v_1.2 in the spinal cord white matter also starting at 7 dpi(*44*). For this purpose, adult female rats were subjected to a force-controlled spinal contusion at T10 after laminectomy and were assessed at various timepoints after injury via [^18^F]3F4AP PET imaging, followed by *ex vivo* evaluation of the spinal cord tissue by autoradiography, histological staining for myelin content and immunohistochemical staining of myelin and axonal markers (**Fig 1A**). Behavioral assessment using the Basso-Beattie-Bresnahan (BBB) locomotor scale(*47*) was performed immediately after injury and for 28 days to score the severity of the injury and its corresponding clinical presentation. Immediately after the lesion, rats presented a sharp decrease in locomotor performance (BBB score) at 1 dpi that spontaneously improved until reaching a plateau 2 weeks after injury (**Fig 1B**). This result aligns with previously observed symptomatic demonstration in rats after a moderate severity spinal contusion injury, where behavioral, anatomical (electron microscopy), and electrophysiological (postinjury conduction) assessments showed a marked pattern of demyelination at 1 week after injury(*46*). We therefore pondered if [^18^F]3F4AP could be used to detect spared demyelinated fibers around the peak of disease and after the initial inflammatory response has subsided in the subacute injury phase.

**Fig. 1.**
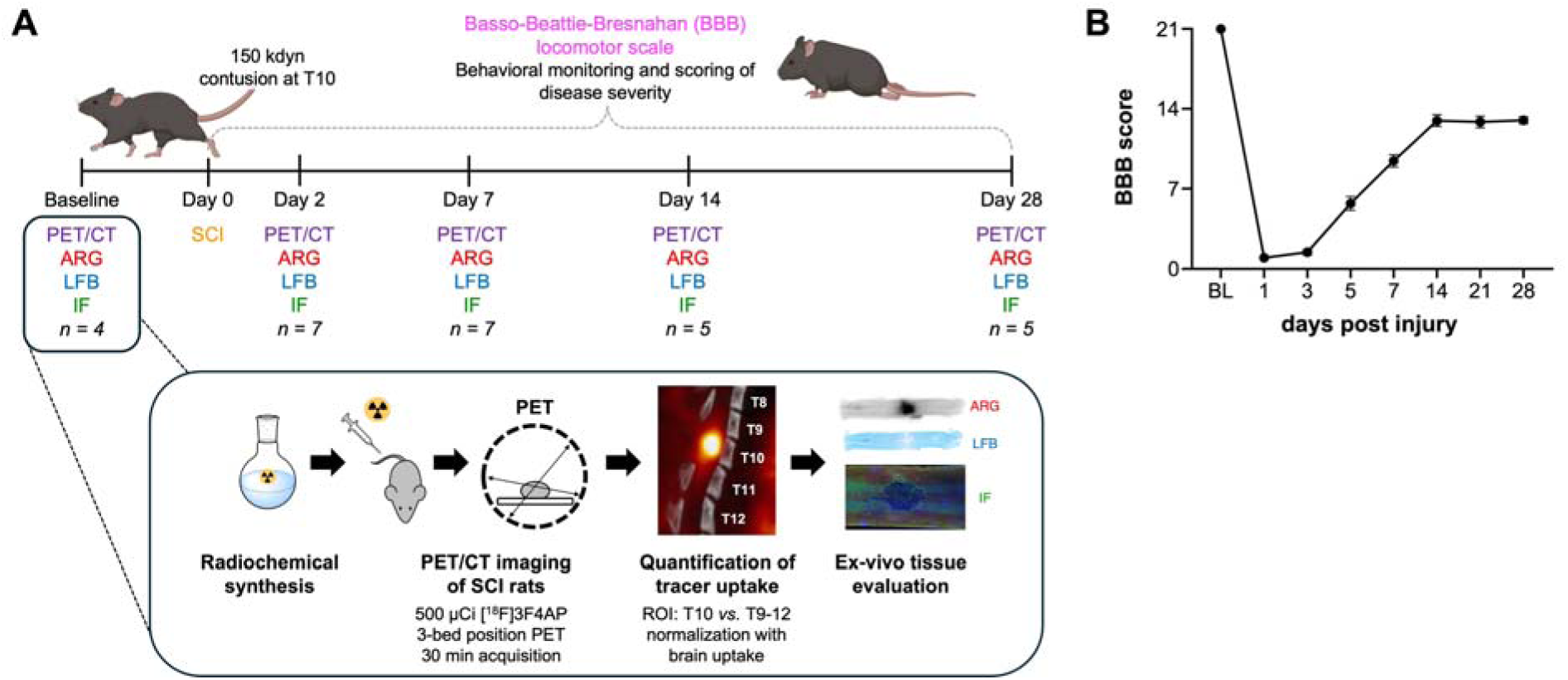
Cross-sectional evaluation of rat SCI with [^18^F]3F4AP. (A) Rats were subjected to a spinal contusion injury and evaluated at different timepoints (baseline, 2, 7, 14 and 28 days-post-injury (dpi)) with [^18^F]3F4AP PET and post-mortem evaluation of the spinal cord tissue at each timepoint. (B) Clinical evaluation was performed at baseline (BL) and at 1, 3, 5, 7, 14, 21, and 28 dpi using the BBB score.

### Imaging demyelinated axons in rats 7 days after spinal contusion

Based on previous reports of peak demyelination and conduction loss 7 days after spinal contusion(*46*), we chose to initially evaluate [^18^F]3F4AP binding in the injury at this timepoint. To assess tracer binding, rats were scanned in supine position to minimize motion of the spine due to breathing during the scan using a multibed position acquisition protocol consisting of a 0-30 min dynamic scan of the trunk region, a 5 min static acquisition of the head, a 5 min static acquisition of the pelvic area, followed by a CT scan for anatomical reference. On the whole-body PET/CT images, there is an area of high focal uptake at the site of injury (**Fig 2A**). Magnification of the coregistered PET and CT images shows that the area of high focal uptake corresponds to the location of the impact beneath the laminectomy (T10) (**Fig. 2B**). Following the scan, one animal was euthanized and their cord extruded for autoradiography, confirming an area of very high focal uptake within the impacted cord (**Fig. 2C**). Time-activity curves (TACs) of the injury (T10) and adjacent spinal cord segments (T8-T12) show slower washout of the tracer at T10 than the other vertebral segments, indicative of greater tracer binding (**Fig. 2D**). Based on the TACs, we selected the standardized uptake value (SUV) from 10-30 min as a measure of binding and quantified the signal at the different spinal cord segments. The mean SUV_10-30 min_ at T10 was 1.12 ± 0.15 (n = 7), which represents a 185 % increase when compared to the mean SUV above the injury at T8 (0.60 ± 0.06) (**Fig 2E**). For further comparisons across animals, values were normalized to brain SUV_30-35 min_ as an internal control. Relative to the brain, T8 showed an SUV ratio (SUVr) of 1.26 ± 0.04 and the lesion showed a SUVr of 2.49 ± 0.09 (n = 7) (**Fig 2F**). Notably, the high uptake at the injury could also be clearly observed in a standard clinical PET/CT scanner. A subset of rats that was euthanized 30 min after tracer administration and scanned in a clinical scanner displayed a distinct and focal uptake in the injured area characteristic of the lesion site (**Fig. S1**).

**Fig 2.**
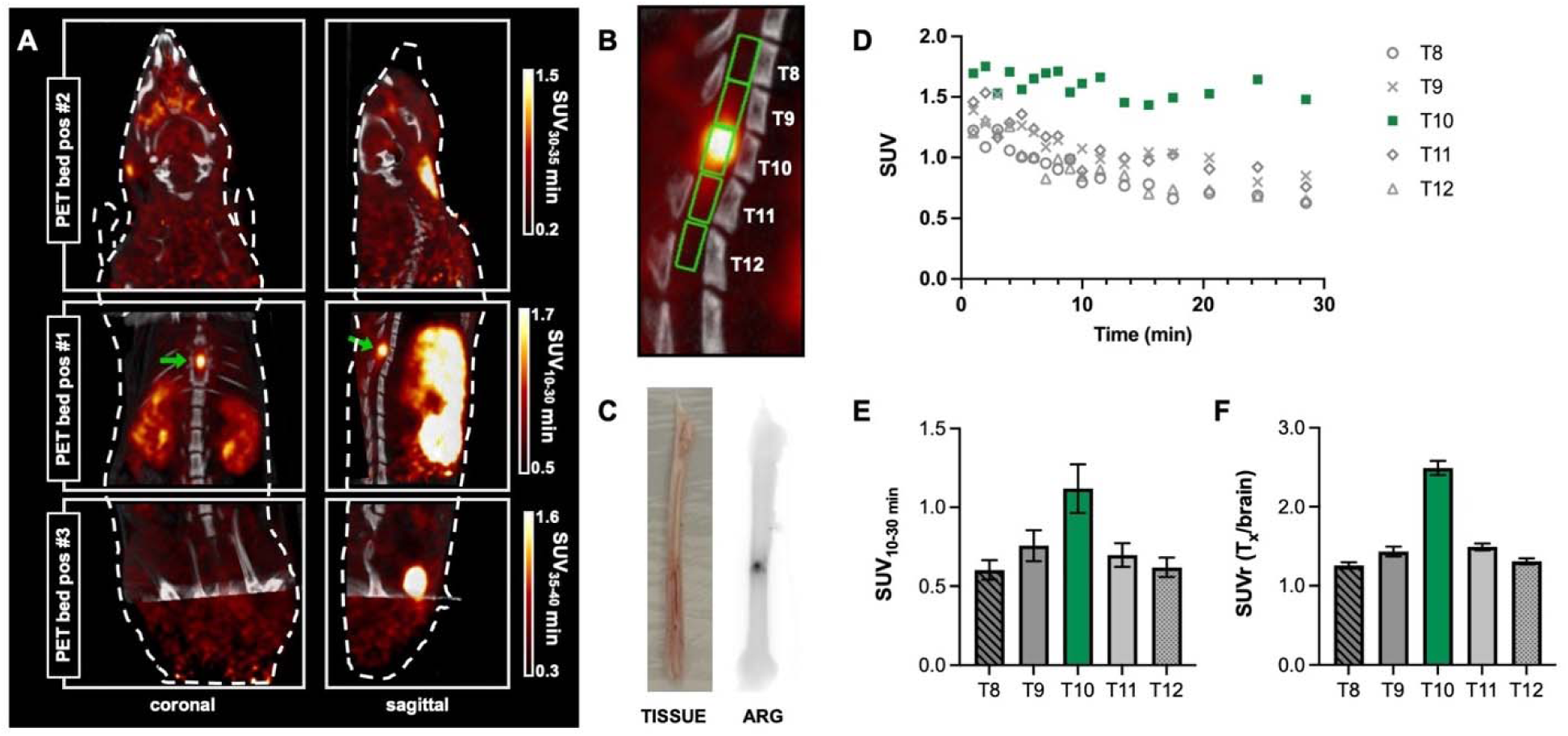
Evaluation of [^18^F]3F4AP in rodent spinal cord injury. (A) Representative coronal and sagittal views of whole body (3-bed position) [^18^F]3F4AP PET/CT in a rat at 7 dpi. (B) Magnification of PET/CT images showing laminectomy at T10, high PET signal in the injured cord and regions of interest (ROIs) selected for quantification. (C) Explanted spinal cord (left) and corresponding autoradiography (ARG) (right) of one rat. (D) Representative time-activity curves (TACs) extracted from the ROIs corresponding to lesion site (T10, green) and surrounding vertebral segments (T8-T12). Quantification of (E) vertebral ROI SUV from 10 to 30 min and (F) normalized vertebral SUV to brain uptake (n = 7). Data are mean ± SEM. Representative PET images and *ex-vivo* tissue correspond to different animals.

### Imaging injury progression at different timepoints after spinal contusion

After observing high [^18^F]3F4AP uptake at the injury at 7 dpi, we examined if [^18^F]3F4AP imaging could be used to monitor injury progression at different timepoints after contusion. For this purpose, we designed a cross-sectional study, where PET imaging was performed in distinct subsets of animals at a specific timepoint (baseline, 2, 7, 14 and 28 dpi; **Fig 3A**) and followed by *ex vivo* evaluation of the post-mortem spinal cord tissue. Increased uptake at the injury site (T10) when compared to surrounding cord segments was observed at 7, 14 and 28 dpi, but not at 2 dpi, (SUVr = 1.14 ± 0.10 (baseline, n = 4), 1.44 ± 0.02 (2 dpi, n = 7), 2.49 ± 0.09 (7 dpi, n = 7), 2.26 ± 0.25 (14 dpi, n = 5), 2.57 ± 0.23 (28 dpi, n = 5)) which suggests the presence of demyelinated fibers after the acute injury phase has subsided (**Fig 3B**). Interestingly, no significant differences were observed in SUVr for the lesion epicenter (T10) after the 7 dpi peak, but there is a more pronounced apparent spreading of the injury to contiguous vertebral segments at later timepoints. When comparing SUVr values of surrounding segments T9 and T11 at the 7, 14, and 28 dpi timepoints it is evident that tracer binding continues to increase at these adjacent levels with disease progression whereas the lesion epicenter SUVr values seem to stabilize after the 7 dpi timepoint, which suggests spreading of demyelination (**Fig. S2**). Notably, the stable imaging from day 14 onwards is consistent with the plateau observed in clinical score observed in these animals.

**Fig 3.**
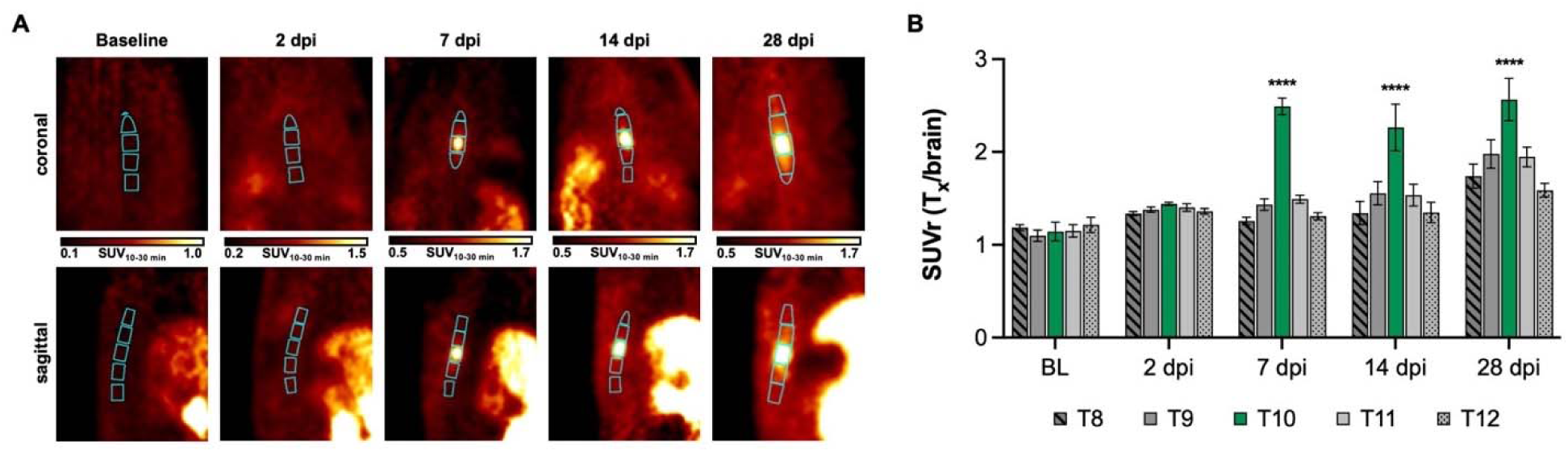
Longitudinal evaluation of [^18^F]3F4AP in rodent SCI. (A) Representative coronal and sagittal views of thoracic [^18^F]3F4AP PET at baseline (BL), 2, 7, 14, and 28 dpi showing CT-based ROI selection for vertebral segments T8-T12. (B) SUVr quantification of SUV_10-30min_ from injury ROI (T10, green) and adjacent vertebral segments (T8-T12) normalized to whole brain SUV_30-35 min_. Statistical analysis was performed using a two-way ANOVA with Dunnett’s multiple comparison test. * denotes comparison versus BL at T10 (\*\*\*\**p* < 0.0001). For *in vivo* PET quantification, BL *n* = 4, 2 dpi *n* = 7, 7 dpi *n* = 7, 14 dpi *n* = 5, and 28 dpi *n* = 5. Data are mean ± SEM.

### *Ex vivo* tissue evaluation correlates with imaging and clinical assessment

To corroborate that the increased PET signal observed after injury is a result of the binding to demyelinated axons and not to other cells or processes, spinal cord tissue was evaluated by *ex vivo* autoradiography and histochemical staining of myelin with Luxol Fast Blue (LFB) (**Fig 4**). The observed *ex vivo* autoradiographic signal correlated positively with the PET imaging results, where an increase in uptake was detected at the lesion site at 7 dpi and after, but not at 2 dpi. Since disruption of the blood-spinal cord barrier and infiltration of inflammatory cells are expected to peak 1-2 days post injury(*48–50*), the lack of tracer accumulation at 2 dpi strongly suggests that those processes have minimal effect on the [^18^F]3F4AP signal. Furthermore, close examination and comparison of corresponding autoradiography and LFB staining images supported that the increase in tracer uptake originates from demyelinated white matter bands around the lesion, and not from the grey matter or the cavity. This concurs with binding to demyelinated axons and is inconsistent with binding to infiltrating inflammatory cells or accumulation at the cavity due to changes in blood flow (**Fig 4A**). Additionally, it is evident that at later time points (14 and 28 dpi) the area of demyelinated fibers extends further from the cavity, indicative of the more diffuse injury observed in the PET imaging (**Fig 4B**). Spinal cord tissue from a separate cohort of animals was further assessed for reactive astrocyte, axonal and myelin content with immunofluorescence staining of GFAP (glial fibrillary acidic protein), NF200 (neurofilament 200) and MBP (myelin basic protein), respectively (**Fig 4C** and **4D**). Comparison of intact and injured cord showed a higher density of reactive astrocytes near the injury, as well as greater occurrence of demyelinated fibers near the lesion.

**Fig 4.**
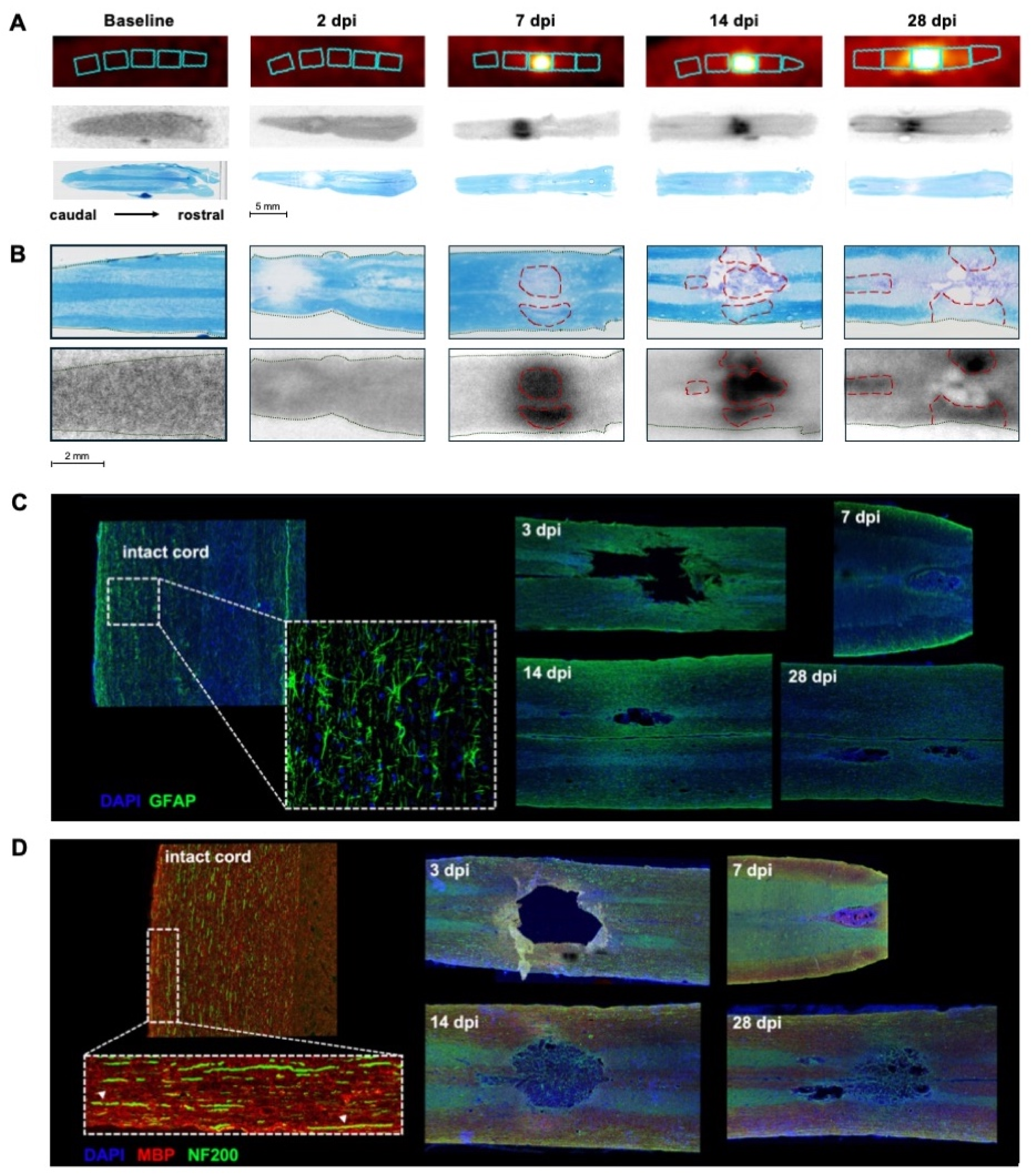
Evaluation of ex-vivo SCI tissue. A) Corresponding PET imaging, autoradiography, and Luxol Fast Blue (LFB) myelin staining of rat spinal cord segments for each timepoint. B) Higher magnification autoradiography and LFB images showing tracer uptake and myelin staining in white matter tracts and injury spreading. C) Representative immunofluorescence staining of intact and injured spinal cord at 3, 7, 14 and 28 dpi with DAPI (nuclear, blue) and GFAP (reactive astrocytes, green). D) Representative immunofluorescence staining of intact and injured spinal cord at 3, 7, 14 and 28 dpi with DAPI (nuclear, blue), MBP (myelin basic protein, red) and NF200 (axonal marker, green). For immunofluorescence staining, n = 1 per timepoint.

To further confirm that the binding of the tracer is specific to K^+^ channels in the injury, *in vitro* autoradiography was performed on the same fresh-frozen unfixed tissue sections used for *ex vivo* autoradiography. As it can be seen in **Figure 5**, tracer binding at the injury was only observed when the tracer was administered to the live animal (*ex vivo* autoradiography) and not when the tracer was applied to the tissue sections (*in vitro* autoradiography). Tracer application to tissue sections produced minimal to no binding at the injury or throughout the tissue. Furthermore, addition of excess amount of nonradioactive 3F4AP did not change the result in further decreasing binding. This experiment represents a strong indication of specific binding, as it is known that binding to K^+^ channels requires the channels to be in an open conformation(*51*) that only exists in living tissue when the membrane is depolarized. On the other hand, if the [^18^F]3F4AP signal was nonspecific, it would be expected to remain in postmortem tissue, which did not occur in this case.

**Fig. 5.**
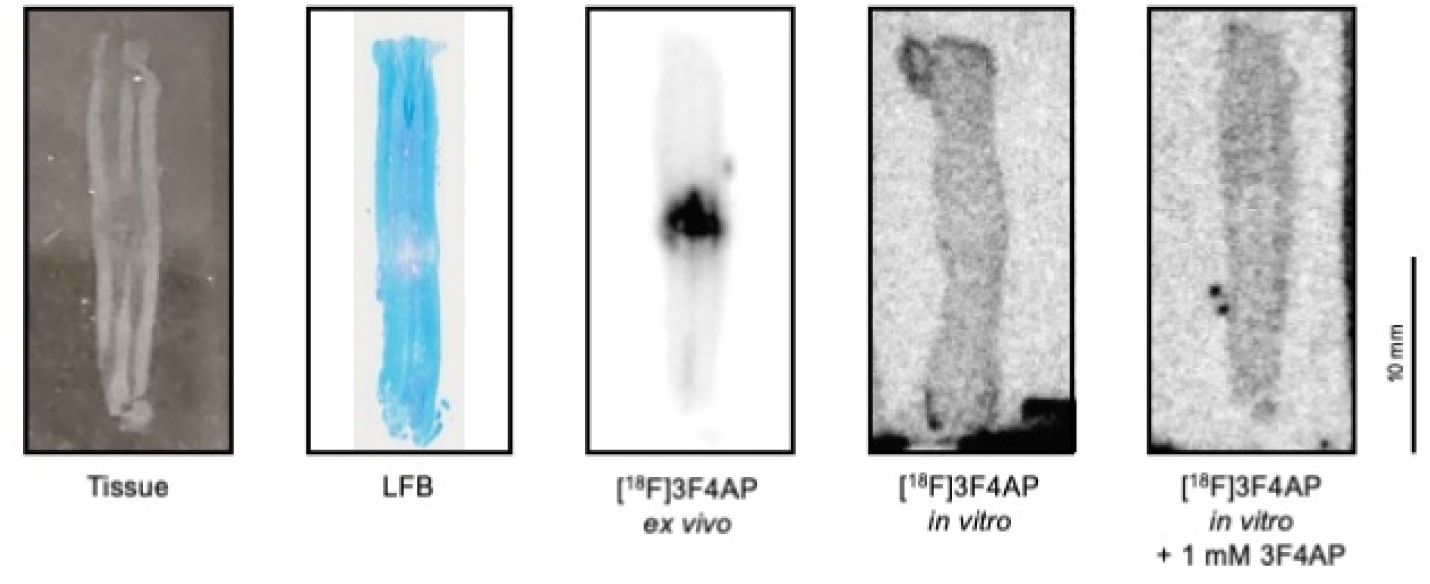
[^18^F]3F4AP specifically binds to exposed K^+^ channels *in vivo*. Representative tissue photograph, LFB staining, *ex vivo* autoradiography, *in vitro* autoradiography, and *in vitro* autoradiography with blocking dose of 3F4AP. Specific binding in the lesion is only observed when the tracer is administered to the live animal followed by tissue dissection and slicing (*ex vivo* autoradiography). Incubation of tissue sections with [^18^F]3F4AP in the absence or presence of nonradioactive 3F4AP results in no binding indicating that the binding at the lesion *in vivo* is specific.

### [^18^F]3F4AP imaging in two human subjects after spinal cord injury

Motivated by the high sensitivity of [^18^F]3F4AP PET imaging in SCI rats, we decided to pursue a pilot study in humans after spinal cord injury. Two subjects with injuries of different severity and etiologies were recruited. The characteristics of the subjects and their injuries are shown in Table 1.

**Table 1.** Characteristics of human SCI volunteers.

| Subject | Sex | Location | Severity | Interval between injury and scan | Etiology |
| --- | --- | --- | --- | --- | --- |
| SCI01 | Male | T12 | AIS-C (incomplete) | 2 years, 5 months | Fall with traumatic burst fracture |
| SCI02 | Male | T11 | AIS-D (incomplete) | 7 years, 5 months | Ruptured spinal cavernous hemangioma |

To evaluate the performance of the tracer, subjects underwent [^18^F]3F4AP PET scans of the injury area, preceded by low dose CT scans for anatomical reference and attenuation correction. The PET acquisition protocol consisted of dynamic scans from 0-45 min and 75-106 min. This dual-window acquisition aimed to capture images of the initial tracer delivery, providing insights into blood perfusion to the injured cord, as well as images from tracer binding in the injured cord.

At the time of the scan, subject SCI01 had an incomplete spinal cord injury at T12 (Asia Impairment Scale (AIS) score = C) as a result from a traumatic fall 2.5 years prior to imaging and used a wheelchair. The burst fracture at T12 and the stabilization hardware was visible on the CT (**Fig. 6A**). Early PET images (0-10 min) show the compressed vertebral body and a very pronounced decrease in signal below the injury (**Fig. 6B**). Quantification of the SUV_0-10min_ in the different spinal segments shows a -76% decrease in PET signal below the injury (L2-L4) relative to above the injury (T10-T11) (**Fig. 6E**). This reduction in signal is likely due to reduced blood perfusion as indicated by the lower initial peak on the TACs of different spinal segments (**Fig. S3**). The late PET images (75-106 min) show a moderate reduction in PET signal below the injury (**Fig. 6C**). Quantification of the SUV_75-106min_ shows a -21% reduction below the injury (L2-L4) compared to above the injury (T10-T11), which may reflect reduced tracer binding due to partial axonal loss (**Fig. 6F**).

**Fig. 6.**
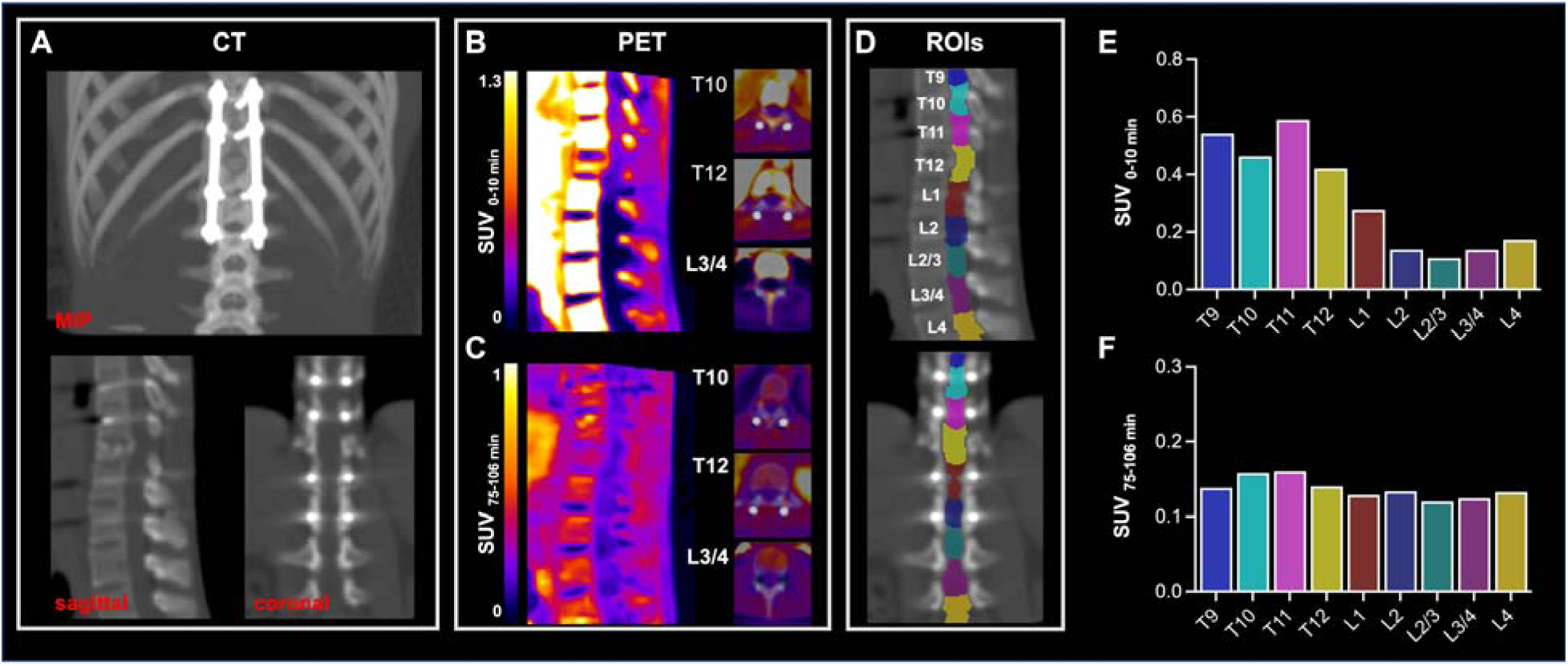
[^18^F]3F4AP in a human subject after severe incomplete spinal cord injury. A) MIP, sagittal and coronal CT images showing location of metal stabilization hardware and crushed vertebral body at T12. B) Early PET sagittal images showing compressed vertebral body and low PET signal in the cord below the injury. C) Late PET sagittal images showing reduced PET signal in the cord below the injury. Axial images at T10, T12 and L3/4 are shown for both early and late PET. D) Selected ROIs for segmentation of the spinal cord. E) Quantification of SUV_0-10 min_ and F) SUV_75-106 min_ at the selected spinal segments.

Subject SCI02 had an incomplete spinal cord injury at T11 (AIS = D) due to a burst spinal hemangioma 7.5 years prior to imaging, which caused initial paralysis below the injury. After T10-T12 laminectomies (visible on CT, **Fig. 7A**) and removal of intramedullary lesion, the subject gradually recovered and by the time of the scan could walk with minor gait disturbances. The subject, who had no metal hardware, also underwent a separate spinal 3T MRI at the injury location. On the MRI, there was T2 hyperintense region near the injury compatible with demyelination (**Fig. 7B**). Early PET images show a localized area of increased tracer signal when compared to spinal cord segments above the injury (+104%) (**Fig. 7C** and **7F**). This increase in signal is suggestive of increased blood perfusion possibly due to ongoing inflammation, as can be seen in the TACs (**Fig. S4**). Late PET images showed +26% increase in signal that colocalized with the T2 hyperintensity consistent with demyelination (**Fig. 7D** and **7G**).

**Fig. 7.**
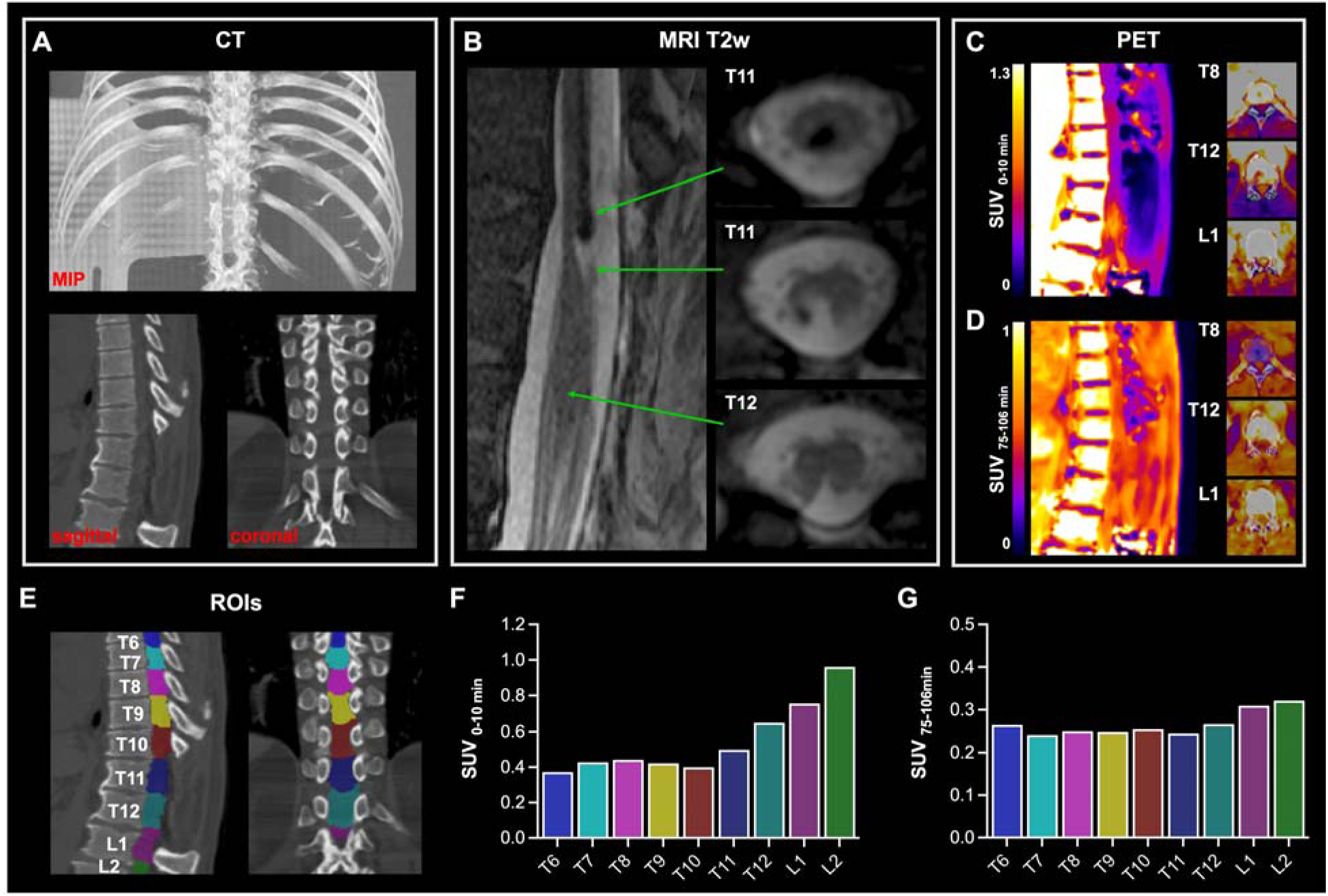
[^18^F]3F4AP in a human subject after mild incomplete spinal cord injury. A) MIP, sagittal and coronal CT images showing laminectomies at T10-T12. B) T2w MR sagittal and axial images showing the spinal lesion. C) Early PET sagittal images showing high PET signal in the cord around injury. D) Late PET sagittal images showing increased PET signal in the cord around the injury. Axial images at T8, T12 and L1 are shown for both early and late PET. E) Selected ROIs for segmentation of the spinal cord. F) Quantification of SUV_0-10 min_ and G) SUV_75-106 min_ at the selected spinal segments.

## DISCUSSION

SCI can result in devastating effects on quality of life. Medical imaging can provide information for accurate disease prognosis and treatment monitoring. Residual tissue bridges and MRI based metrics of myelination at the level of the spinal cord injury can be predictive of subsequent motor recovery(*52*). While CT and MRI are most used in the clinic, PET imaging can offer biochemically specific and quantitative information that can complement these modalities. Furthermore, while MRI can reveal anatomical changes after injury such as cord thinning or demyelination, the presence of metal stabilization hardware can often cause artifacts in standard contrasts and hinder susceptibility-based diffusion imaging. Thus, we have investigated the use of the novel PET tracer, [^18^F]3F4AP, following SCI and describe its sensitivity and potential as a diagnostic tool for assessing the extent of demyelination and axonal damage after injury.

The choice of using [^18^F]3F4AP in SCI was motivated by prior studies that show some effectiveness of 4AP in SCI treatment(*34–42*) and by the increased uptake of [^18^F]3F4AP observed in a traumatic brain injury in a rhesus macaque(*28*). Our results demonstrate that [^18^F]3F4AP exhibits a high sensitivity to incomplete SCI in rats. In this study, we used a spinal contusion model in rats, which results in a severe injury characterized by partial paralysis below the injury immediately after the procedure, followed by a slow but progressive recovery. Previous studies using similar models have shown sparing of axons with severe demyelination and upregulation of axonal K^+^ channels at the injury site starting around 7 days post injury(*43, 44*). The PET findings revealed a more than 2-fold increase in tracer binding, highly localized to the injured segment of the cord at 7 dpi relative to baseline. Notably, the [^18^F]3F4AP signal evolved over time. In the acute SCI phase (2 dpi), there was no discernible increase in tracer uptake compared to baseline, indicating an early phase in which damaged myelin has not yet been removed and axonal K_v_ channels have not yet redistributed. This observed delay in K_v_ channel redistribution and upregulation after injury has been described by Karimi-Abdoolrezaee in a compressive SCI model, where this process does not appear to be an early response to injury but rather more present at more chronic stages of SCI. This finding also supports that [^18^F]3F4AP signal is not dependent on blood-spinal cord barrier damage or inflammation since these processes are expected to peak very soon after the injury. By one week post injury, the signal reached its maximal intensity, aligning with the peak of demyelination. Subsequently, at 14 and 28 days, the signal began to spread out, consistent with the progression of demyelination.

The agreement between our PET findings and autoradiography, coupled with LFB staining for myelin, supports the notion that the [^18^F]3F4AP signal primarily originates from demyelinated axons. Close up views of paired sections imaged by *ex vivo* autoradiography and LFB staining clearly show that [^18^F]3F4AP primarily localizes to demyelinated white matter bands and not to the cavity or grey matter. Representative immunohistochemistry imaging of one animal at each point corroborated a high concentration of demyelinated axons at day 7, 14 and 28 post injury consistent with the changes observed by PET, autoradiography and LFB. Furthermore, *in vitro* autoradiography on the same tissue sections showed no binding in the injured tissue which strongly suggests that the binding is specific to K^+^ channels since 4AP and derivatives can only bind to open K^+^ channels in living tissue, whereas non-specific binding would likely be present in post-mortem tissue.

Many human spinal cord injuries are incomplete, with some spared circuits that hold potential for recovery of voluntary movement(*53*). Slower and weaker axonal conduction after demyelination has been suggested as a pathophysiological basis for discomplete spinal cord injuries, characterized by apparent complete transection as judged by clinical criteria, but with neurophysiological evidence of conduction through the level of damage(*2, 54, 55*).

Our pilot imaging study in two human subjects after SCI represents a crucial step in the translational potential of [^18^F]3F4AP. The study participant with a burst spinal hemanginoma (SCI02) experienced near-complete recovery, while the study participant with a traumatic fall injury (SCI01) had initial gains in the first month post-injury form AIS-A to AIS-C, but muscle strength and sensation below the injury remained low. Thus, these two participants represent a broad range in terms of the potential to have residual axons crossing the spinal lesion. The subject with the milder injury showed increased binding at and immediately below the injury, suggestive of presence of demyelinated fibers. The smaller increase in tracer binding in this subject compared to the rat SCI injury model may reflect limited demyelination in this particular subject given the nature of the injury and the extended period (7.5 years) between injury and scan. Conversely, the subject with the more severe injury exhibited a decrease in binding at and below the injury site. While these initial observations are promising, further studies with larger cohorts are warranted to validate and generalize these trends.

This study has several limitations that should be considered. Firstly, there were differences in the PET acquisition protocols between humans and rats (*i.e.*, injection on the scanner *vs*. injection on the bench) which allowed to look at tracer delivery in humans but not in rats. Additionally, the timing between injury and imaging was inconsistent, ranging from 2 to 28 days in rats and 2.5 and 7.5 years in humans. While SCI is more prevalent in men than in women, we utilized only female rats in the study due to the easier management of bladder dysfunction after injury. Finally, while previous studies have documented K_v_ channel changes in similar models, in this study we did not directly measure these changes in animal or human tissue samples by immunohistochemistry.

Given the heterogeneity of SCI, a tracer capable of detecting spared axons is of paramount importance. Our study underscores the potential value of [^18^F]3F4AP in capturing this heterogeneity by sensitively detecting demyelination and axonal damage, providing valuable insights into the underlying pathology. Furthermore, while some clinical studies have shown the benefits of using 4AP in SCI patients(*34–39*), others have not(*40–42*), raising the question of whether demyelination plays a role in many SCI cases and highlighting the need for a biomarker that can predict response to 4AP.

In conclusion, our study represents the first investigation of [^18^F]3F4AP in traumatic spinal cord injuries, demonstrating its high sensitivity and specificity for demyelinated axons after SCI. As we move forward, further studies with larger cohorts and longitudinal assessments are crucial to establish the reliability of [^18^F]3F4AP PET as a diagnostic tool. Additionally, exploring its potential for monitoring therapeutic interventions and its broader applicability in diverse SCI populations will be essential for clinical translation. The promising results presented here lay the groundwork for future investigations into the clinical utility of [^18^F]3F4AP in the realm of spinal cord injury.

## MATERIALS AND METHODS

### Study Design

Here, we sought to investigate the potential of the PET tracer [^18^F]3F4AP to image demyelination after spinal cord injuries (SCIs). A rat model of incomplete SCI with a well-established time course of demyelination was selected to assess the temporal changes in tracer uptake. Spinal cord tissue was evaluated after PET scanning by *ex vivo* autoradiography and histological staining of myelin to correlate imaging results. Tissue from a separate cohort of rats (one animal per timepoint) was collected and evaluated by immunofluorescence staining for qualitative comparison. A total of 35 rats (female Sprague Dawley, Charles River Laboratory) were used for this work. Sample size was determined empirically to satisfy the field standards and have sufficient power to detect statistical differences. Experiments were not blinded. To assess the potential translatability of the findings to humans, two human subjects with SCI of different etiology and duration were also scanned with the tracer and the images compared with CT and MRI.

All rodent procedures were approved by the Institutional Animal Care and Use Committee (IACUC) at the Massachusetts General Hospital. All animal studies were conducted in compliance with the ARRIVE guidelines (Animal Research: Reporting in Vivo Experiments) for reporting animal experiments.

Human imaging studies were performed in line with the principles of the Declaration of Helsinki. Approval was granted by the Institutional Review Board at the Massachusetts General Hospital (IRB# 2020P003898, PI: Linnman). [^18^F]3F4AP was administered under an investigational new drug (IND) authorization from the U.S. Food and Drug Administration (FDA) (IND # 135,532, Sponsor: Brugarolas).

### Animal subjects

Adult female Sprague Dawley rats (*n* = 28, 8-12 weeks-old) were subjected to a clinically relevant SCI model in combination with microPET studies, *ex-vivo* examination of the spinal cord tissue and behavioral assessment to evaluate the temporal pattern of demyelination. Moderate severity spinal contusion injury was performed as previously described(*46*). Briefly, rats were anesthetized with isoflurane, laminectomy at the mid-thoracic level was performed and a contusion injury was delivered through the intact dura using a force-controlled spinal cord impactor (IH-0400, Infinite Horizon) set to 150 kdyn. At various postinjury timepoints, animals were removed for PET imaging, autoradiographic and histological studies (baseline: *n* = 4, 2 dpi: *n* = 7, 7 dpi: *n* = 7, 14 dpi: *n* = 5, 28 dpi: *n* = 5). Clinical evaluation was performed throughout the study.

### Clinical evaluation of SCI rats

Basso, Beattie, and Bresnahan (BBB) locomotor rating scale was used to evaluate locomotor performance at baseline, and 1, 3, 5, 7, 14, 21 and 28 days after the lesion. Rats were placed in an open field arena and both hindlimbs were assessed during spontaneous locomotion over a 4-minute session. Scores were calculated according to the 22-point BBB scale for each hindlimb and then averaged for each rat.

### Human subjects

Two male volunteers participated in the study after providing informed consent. Participating criteria included adults between 18 and 65 years old with a history of spinal cord injury and no contraindications to the study such as severe claustrophobia, inability to provide inform consent or accumulated annual radiation dose greater than 30 mSv. No subjects were excluded on the bases of sex, ethnicity, or race.

### Radiotracer production for animal studies

[^18^F]3F4AP was produced in a GE TRACERlab Fx2N synthesizer according to previously reported procedure(*56*). Semipreparative HPLC separations were performed on Sykam S1122 Solvent Delivery System HPLC pump with the UV detector at 254 nm with a Waters C18 preparative column (XBridge BEH C18 OBD Prep Column 130 Å, 5 µm, 10 mm × 250 mm). The obtained [^18^F]3F4AP fraction (radiochemical purity > 99%, molar activity = 3.0 ± 0.9 Ci/µmol) at end of synthesis (EoS); n = 7), in 95% 20 mM sodium phosphate buffer, pH 8.0, 5% EtOH solution) passed the QC (with Thermo Scientific Dionex Ultimate 3000 UHPLC equipped with Waters XBridge BEH C18 analytical column [130 Å, 3.5 µm, 4.6 × 100 mm], 95% 10 mM sodium phosphate buffer, pH 8.0, 5% EtOH solution as mobile phase, t_R_ = 4.7 min) and was ready for injection into animals.

### Radiotracer production for human use

[^18^F]3F4AP was produced by the MGH Gordon PET Core cGMP radiopharmacy using a Neptis ORA synthesizer as previously communicated(*57*). The synthesis method is based on the previous report by Basuli *et al*(*58*). The tracer was purified using a semiprep HPLC column (Waters XBridge C-18, 5 μm, 10 × 250 mm) using 20 mM sodium phosphate (pH 8) mobile phase containing 5% ethanol at a flow rate of 4 mL/min. The HPLC fraction containing the product (approx. t_R_ 10-11 min) was diluted with 10 mL of 0.9% sodium chloride for injection, USP, and passed through a 0.22 μm sterilizing PES filter into a vented 30 mL sterile empty vial. The product vial was visually inspected and quality control was performed to FDA and USP standards for chemical identity and purity, radiochemical purity, pH, radionuclidic identity, residual solvents, sterility, and bacterial endotoxins. Identity was confirmed by coinjection of authentic standard on an analytical HPLC column (Phenomenex Gemini C-18, 5 μm, 4.6 × 250 mm) with mobile phase 10 mM sodium phosphate dibasic, 0.25% triethylamine and 5% acetonitrile at a flow rate of 1 mL/min (t_R_ = 10 min). Purity was assessed by area under the curve (AUC) of the product peak at 239 nm relative to other peaks not present in the blank. Molar activity (MA) at the time of injection was 7.59 Ci/µmol (SCI01) and 3.68 Ci/µmol (SCI02). The sterile filter used was tested for integrity. The dose was released for injection after passing all quality control tests except for sterility, which was performed after release. All the batches of [^18^F]3F4AP used in this study met all product specifications, including sterility test.

### Animal PET/CT image acquisition and reconstruction

Rats were anesthetized with isoflurane gas and intravenously administered 500 µCi [^18^F]3F4AP. Immediately after injection, rats were set in supine position and multi-bed position PET was acquired in a tri-modality PET/SPECT/CT scanner (Triumph LabPET from Trifoil imaging) as follows: bed position #1 – dynamic PET for 30 min post tracer administration centered at the thoracic injury site, bed position #2 – static acquisition for 5 min at the head, and bed position #3 – static acquisition for 5 min at the pelvic area. CT was acquired in 2-bed position following PET for anatomical reference.

### Animal PET/CT image analysis

PET images were analyzed and visualized using AMIDE. PET and CT were coregistered. Using CT for anatomical reference, laminectomy site was located and cylindric regions of interest (ROIs) were drawn for vertebral segments from T9 to T12. PET summed uptake values from 10-30 min after injection were extracted and normalized to brain uptake values from 30-35 min.

### Dissection and ex-vivo autoradiography

Immediately following the completion of the PET/CT scan, the rats were euthanized by intraperitoneal injection of pentobarbital sodium and phenytoin sodium solution (Euthasol^®^) and the spinal cord between approximately T1 and L5 isolated by hydraulic extrusion into ice cold PBS (similar to PMID: 28190031). The spinal cord was patted dry on a paper towel and embedded in OCT in a cryostat mold. Next the spinal cord was sliced longitudinally at 20 µm thick sections using a cryostat, mounted onto microscope slides, and allowed to air dry. The slides were apposed onto a Phosporimager screen overnight. The next day, the screens were imaged using a Typhoon FLA 9000 Phosphorimager.

### In vitro autoradiography

Selected tissue sections on microscope slides were incubated with 5 nCi of [^18^F]3F4AP inside a moisture chamber for 30 min. Afterwards, the solution containing [^18^F]3F4AP was discarded and the tissue sections were washed 3 times with ice cold PBS for 30 seconds. The slides were air dried, apposed to a Phosphorimager screen overnight and imaged using a Phosphorimager the next day.

### Luxol fast blue staining

For Luxol fast blue (LFB) staining, selected frozen tissue sections on microscope slides corresponding to previous autoradiography slides were stained with a standard staining protocol. Briefly, slides were incubated in 0.1% LFB in acidified 95% ethanol overnight at 56 °C. The slides were then differentiated and counterstained with 0.05% lithium carbonate and 0.1% cresyl violet solution.

### Histology and microscopy

Anesthetized rats (*n* = 4) were transcardially perfused with 4% paraformaldehyde (PFA). Dissected spinal cords were post-fixed in 4% PFA overnight and infiltrated with 30% sucrose solution for cryoprotection before sectioning. 40 µm thick sections were cut and kept free floating in PBS at 4 °C. For immunohistochemistry, tissue was blocked for one hour in PBS with 10% horse serum and 0.3% Triton X-100, and then incubated overnight at 4 °C in primary antibodies diluted in PBS with 10 % horse serum and 0. 3% Triton X-100, followed by incubation of secondary antibodies (Invitrogen donkey anti-rabbit Alexa Fluor 488, donkey anti-rat Alexa Fluor 555) for 4 hr at room temperature after washing 3 times for 15 minutes. Primary antibodies used were rabbit anti-GFAP (1:1000, Dako Z0334); rat anti-MBP (1:2000, Abcam ab7349); rabbit anti-NF200 (1:400, Sigma N4142). After staining, free floating tissue was washed, adhered to microscope slides, and mounted with Fluoromount-G™ Mounting Medium, with DAPI (Invitrogen). Fluorescent images were acquired using a Zeiss LSM900 Confocal Microscope and a Leica Stellaris Confocal Microscope. Brightness and contrast of the images were adjusted with imageJ (NIH).

### Radiotracer administration to humans

Dose was drawn into a syringe, measured, and administered intravenously as a 1 min infusion through a catheter in the hand or arm. After administration of the dose, the catheter was flushed with 10 mL of saline and the residual activity in the syringe and catheter measured to calculate the injected dose (ID).

### Human PET/CT image acquisition and reconstruction

Imaging was performed on a GE Discovery MI PET/CT scanner. Subjects were positioned on the bed of the scanner and a low dose CT of the injury location was acquired. Dynamic PET images were acquired from 0-45 min and again from 75-106 min. Subjects were encouraged to void their bladder to eliminate radioactive urine during the break. After completion of the scan, the PET data was reconstructed using the scanner’s VUE Point HD reconstruction algorithm with 17 subsets and 3 iterations with corrections for scatter, attenuation, deadtime, random coincidence and scanner normalization. To improve PET image resolution and reduce PET image noise, we adopted the kernel method-based image reconstruction method to generate PET images from raw PET listmode data(*59*). For the kernel method the PET image was represented as a function of features derived from the anatomical image. In our study, the anatomical image came from the CT image during the PET/CT scan. The features of each voxel were the intensity values from the CT image in a patch centered around the considered voxel. Transformation of the extracted features was computed for each voxel and saved as a kernel matrix that was included in the PET list-mode reconstruction framework(*60*). The kernel method-based reconstruction was utilized to generate an early-phase image using 0-10 min of PET data and a late-phase image using the data acquired between 75-106 min.

### Human MRI acquisition and processing

Subject SCI02 underwent MRI on a Siemens 3T MMR scanner utilizing the body coils to visualize the lesion, which was most clearly observed on a sagittal T2 spin echo sequence with a 0.3 mm in-plane resolution and 2 mm slices (flip angle 150°, echo time 109 ms, repetition time 4660 ms), and on a transversal T2 spin echo sequence with 0.6 mm in-plane resolution and 3.9 mm slices (flip angle 150°, echo time 82 ms, repetition time 3500 ms).

### Human PET/CT and MRI image analysis

PET images were analyzed and visualized using VivoQuant. MRI was coregistered to the PET/CT images. Using CT for anatomical reference, the visible portion of the spinal cord was segmented using an automated thresholding algorithm into multiple ROIs approximately corresponding to vertebral segments. ROIs were visually inspected and manually edited. PET time activity curves corresponding to 0-45 min were extracted for each ROI. Axial MR slices corresponding to the ROIs were inspected for changes in T2 intensity.

### Statistical analysis

Statistical analysis of *in vivo* PET was performed using GraphPad Prism (version 10.2). Descriptive statistics including mean, standard deviation (SD) and standard error (SE) were calculated for each group. Two-group *t*-tests and multi-group *t*-tests (e.g., ANOVA) with a significance level of α = 0.05 were used to assess differences among groups. Grouped data are reported as mean ± SEM.

## Acknowledgments

We thank the two study participants, David Lee and Kyle Stewart at the MGH Gordon PET cyclotron facility for producing fluorine-18, Tricia Lacefield and the MGH CCM animal facility staff for rodent transfer and handling, and Nicole DaSilva and the Gordon nuclear medicine technologist staff for assistance in human subject studies.

## Author contributions

Conceptualization: KMRT, PB

Methodology: KMRT, AT, PB

Investigation: KMRT, SC, YPZ, CC, KT, MH, MW, SE, CL

Visualization: KMRT, SC, AT, YS, CL, PB

Formal Analysis: KMRT, PB

Data Curation: KMRT, JB, CL, PB

Resources: PB, CL, ZH

Software: AT, KG

Validation: KMRT

Funding acquisition: PB, ZH, CL

Project administration: KMRT, PB, ZH, CL

Supervision: PB, ZH, CL, KG

Writing – original draft: KMRT, PB

Writing – review & editing: ALL

## Data and materials availability

The datasets generated and/or analysed during the current study are available from the corresponding author upon reasonable request.

