## Supplemental Information for "Imaging demyelinated axons after spinal cord injuries with PET tracer [^18^F]3F4AP"

55 Fruit Street

Bulfinch 051

Boston, MA 02114

TABLE OF CONTENTS

| Page 3 | Supplemental Methods |
| --- | --- |
| Page 4 | Supplemental Figures |
| Page 4 | Figure S1. [^18^F]3F4AP PET imaging of rat SCI in a clinical scanner |
| Page 5 | Figure S2. Spatial and temporal comparisons in [^18^F]3F4AP PET signal |
| Page 6 | Figure S3. Time-activity curves of spinal segments in human subject SCI01 (T12 AIS-C incomplete injury) |
| Page 7 | Figure S4. Time-activity curves of spinal segments in human subject SCI02 (T11 AIS-D incomplete injury) |

**SUPPLEMENTAL METHODS**

*Whole body imaging of rats with spinal cord injuries in a clinical PET/CT scanner*: 7 days after spinal contusion injuries five rats were anesthetized with isoflurane gas and intravenously administered 500 µCi [^18^F]3F4AP. Animals were kept anesthetized until euthanasia by intraperitoneal injection of pentobarbital sodium and phenytoin sodium solution at 30 min after tracer administration. Immediately after euthanasia, rats were placed in prone position inside a closed container and placed inside the bore of a clinical PET/CT scanner (GE Discovery MI PET/CT). A CT was acquired for anatomical reference. Following CT, a whole-body static PET acquisition was done for 15 min. PET images were reconstructed using the Q.Clear algorithm from the scanner manufacturer. Images were visualized using Vivoquant software (version 3.5, inviCRO).

**SUPPLEMENTAL FIGURES**

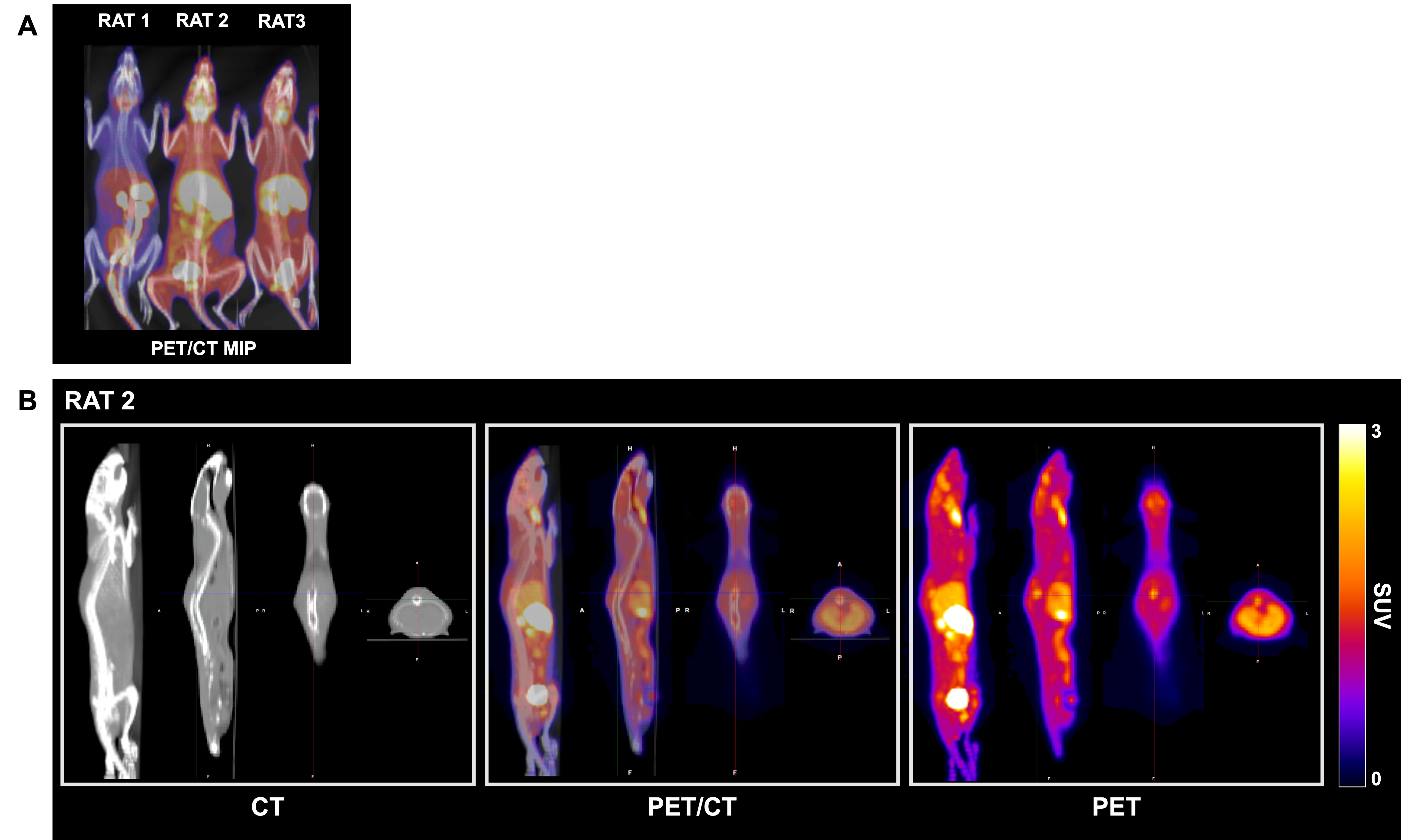

**Figure S1. [^18^F]3F4AP PET imaging of rat SCI in a clinical scanner.** (A) Whole body MIP PET/CT of three rats 7 days post SCI. (B) Multiple views CT, PET/CT and PET images of one rat showing high signal at the injury.

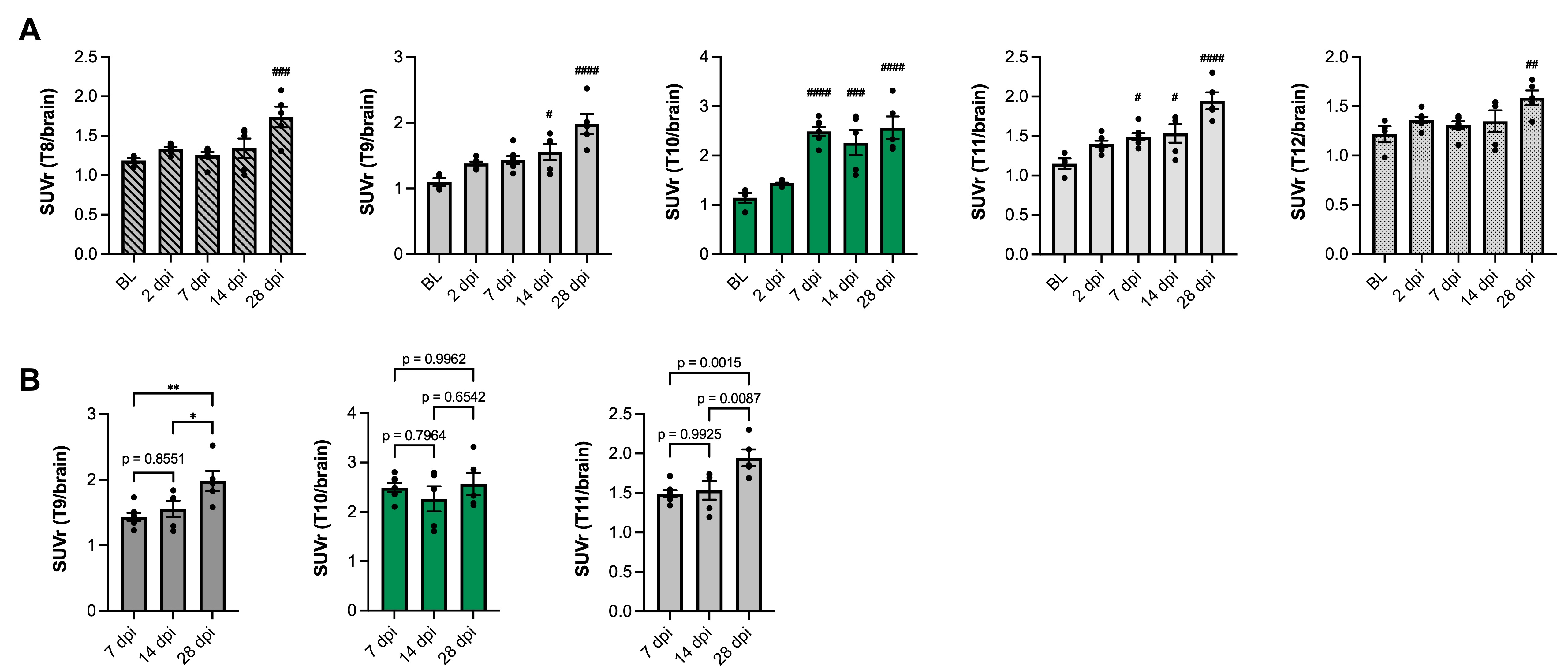

**Figure S2. Spatial and temporal comparisons in [^18^F]3F4AP PET signal.** (A) Comparison in SUVr (SUV_10-30 min_ from vertebral ROI normalized to whole brain SUV_30-35 min_) per spinal segment (T8-12) at each timepoint (baseline (BL), 2, 7, 14, and 28 dpi). Statistical analysis was performed using a one-way ANOVA with Dunnett’s multiple comparison test. # denotes comparison versus BL (^#^*p* < 0.05, ^##^*p* < 0.01, ^###^*p* < 0.001 ^####^*p* < 0.0001). (B) Comparison in SUVr of adjacent (T9, T10) and injury (T10) spinal segments at later timepoints (7, 14, and 28 dpi). Statistical analysis was performed using a one-way ANOVA with Tukey’s multiple comparison test. * denotes direct comparison between groups (^*^*p* = 0.002, ^**^*p* = 0.03). Data are mean ± SEM.

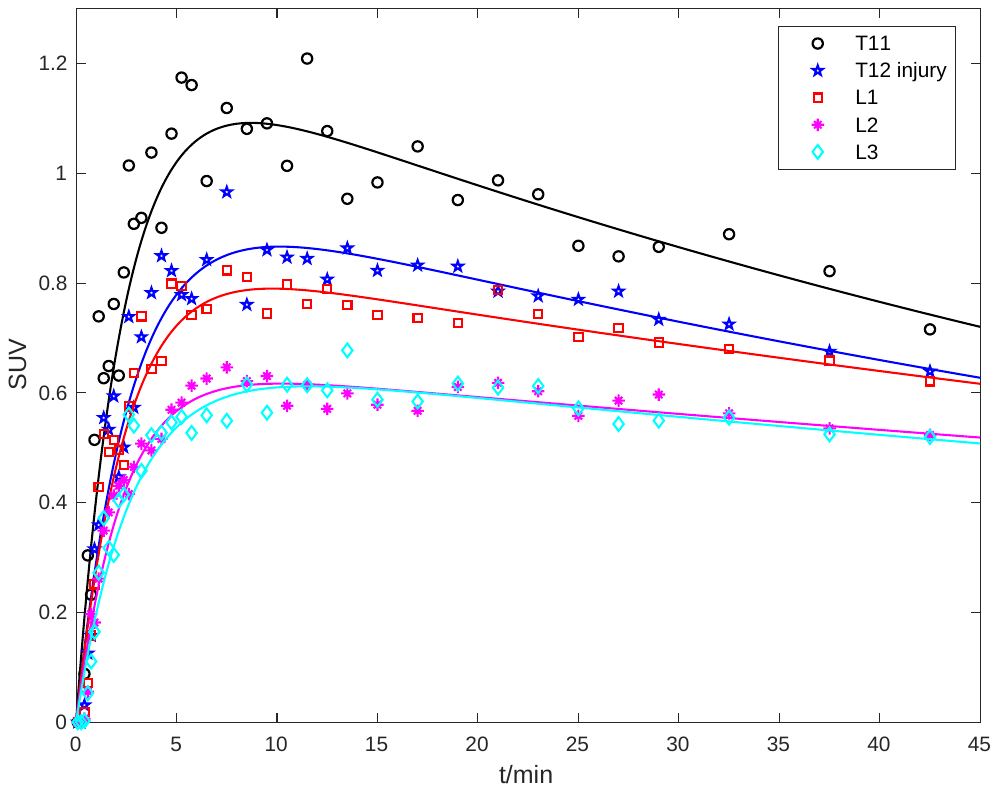

Figure S3. Time-activity curves of spinal segments in human subject SCI01 (T12 AIS-C incomplete injury). There is higher perfusion above the injury (T11) than below the injury (L2 and L3) as evidenced by the height of the peak.

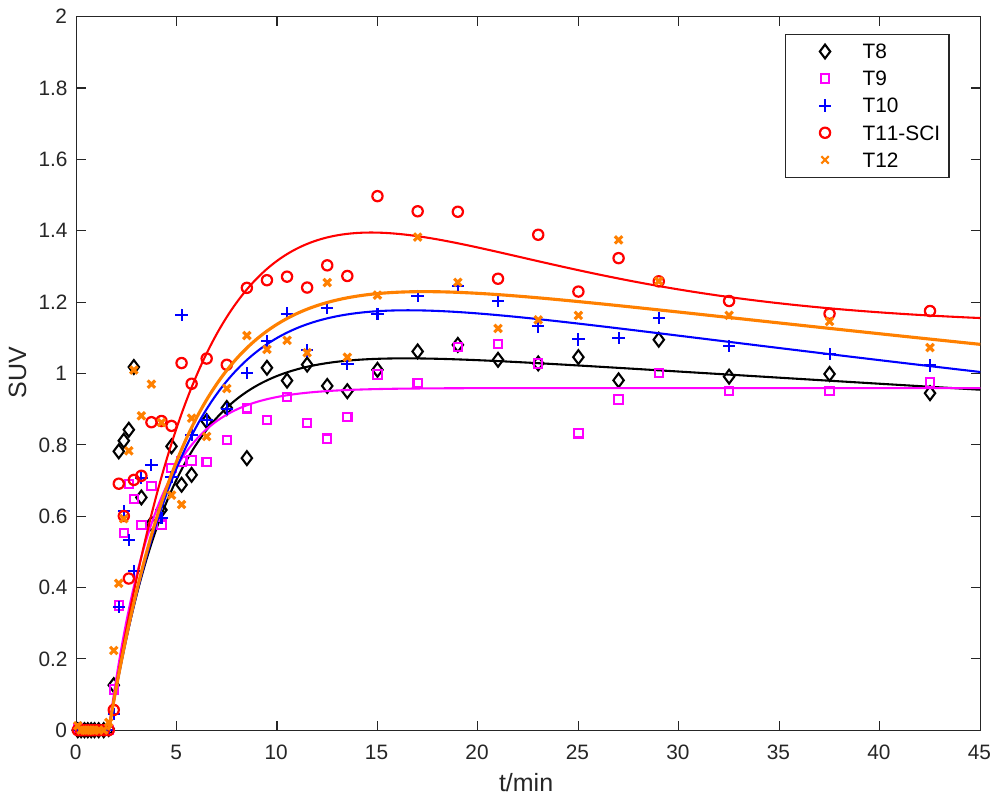

**Figure S4. Time-activity curves of spinal segments in human subject SCI02 (T11 AIS-D incomplete injury).** There is higher perfusion at the injury (T11) relative to other segments evidenced by a higher initial peak.
